# Insights on Skin Adherent Microbiota of the Danube Sturgeon (*Acipenser gueldenstaedtii*)

**DOI:** 10.1101/2024.03.13.584882

**Authors:** Cristian-Emilian Pop, Abdulhusein Jawdhari, Sergiu Fendrihan, Robert Wolff, Nicolai Craciun

## Abstract

The present study assesses the bacterial composition associated with the skin of the Danube Sturgeon, *Acipenser gueldenstaedtii*, emphasizing species that are adherent. Various growth media were employed to cultivate samples collected from a group of 12 sturgeons from the same batch, revealing a diverse and atypical bacterial population. Notably, *Ralstonia pickettii* is documented here for the first time, as there have been no prior associations reported between this species and either wild or aquaculture sturgeon populations, to the best of our knowledge. The identification of adherent and non-adherent bacterial species present on the integument raises questions and concerns regarding the existing microbial communities in sturgeons.

## 1. Introduction

Integumental pathogens pose a growing concern in some sectors of the aquaculture industry, a rapidly expanding food sector of considerable economic significance [1,2]. While the majority of commercially raised fish species intended for human consumption have a rapid growth and high tolerance of bacterial water treatment, mitigating much of the severe impact of pathogens on their population, sturgeons however present a distinct scenario. Due to their prolonged maturation process, that usually takes 8 to 13 years for males and 14 to 23 years for females [3], as well as to their heightened sensitivity to the conventional copper sulphate bacterial treatment [4, 5], the significance of tegument adherent pathogens is therefore accentuated. For sturgeons, and other fish, prolonged pathogenic microbiota hosting can notably impact the growth and survival rate [6, 7] of the fish. In this study we evaluated *Acipenser gueldenstaedtii* skin adherent microbiota, excluding host specific and general parasites [8–10]. In terms of the dermal skeleton, *A. gueldenstaedti* presents five rows of modified diamond-shaped ganoid scales called scutes. These dermal structures are of woven bone type, sharing similarities with the structure of the ewe bone, although the proportion of the apatitic domain seems to be lower for scutes [11]. The outermost layer of the body (covering the scutes as well), is considered tegument, which thus consists of the skin (integument) and its appendices. The skin contains mucosal glands and mucosal skin-associated lymphoid tissues (SALT) [12], which can trigger an innate immune response [13].

The organ (skin) is composed of the outer layer epidermis, followed by dermis and hypodermis. It is the epidermis that possesses a variety of cell types that participate in the exudation of surface mucus [14, 15]. These mucus-producing cells are recognized for harboring diverse biologically active large molecules, primarily glycoproteins, which holds significance in pivotal biological processes, particularly those related to immune responses [16]. The fundamental architectural element of the epidermis is comprised of stratified and irregularly shaped epithelial cells that are organized into multiple layers. These basal layer are composed of columnar basal cells that overlay the basement membrane and such cells possess a columnar nucleus situated toward their upper portion.

The intermediate layer encompasses a variety of multilayered, irregular epithelial cells alongside mucous cells. In the intermediate layer, the cell nuclei are typically round or oval, occupying a central position within the cell. Conversely, epithelial cells in proximity to mucous cells exhibit crescent or irregular nuclei. The interface between the epidermis and dermis is demarcated by a relatively substantial basement membrane [14], the dermis is composed of highly oriented, large and parallel collagenous fibers [15]. In sturgeons, like in many other aquatic organisms, the epidermal mucus represents a line of defense in terms of anti-inflammatory and antimicrobial activity [17].

Still, different skin regions are prone to lesions for various reasons [18], and consequently to infections, offering a gateway for pathogenic and opportunistic bacteria.

Known for being common to many fish species and the aquatic environment in general, the microorganisms identified in this work provide the first insights regarding skin adherent microbiota in the particular case of *A. gueldenstaedti*, although the pattern can be similar for other, if not all sturgeon species.

The purpose of the present work was to identify adherent skin microbiota of cultured *A. gueldenstaedti*, to the best of our knowledge, this is the first such report.

## 2. Materials and Methods

### 2.1. Specimen preparation and sampling

Nine juvenile specimens of *Acipenser gueldenstaedti* (total length ≈ 13 cm) sourced from a fish farming facility were washed through three successive baths of microbiologically pure water, individually rinsed in aseptic conditions, and each placed in an enclosed sterile polycarbonate water tank (AquaMedic, Germany) with 2.5 L of sterile water. A group of three separate specimens were kept as controls (unwashed). Aeration in aseptic conditions was assured with an air pump through sterile silicon tubing fitted with a microbiological air filter. After 8 hours, each specimen was taken out from the designated water tank and rinsed intensively with sterile water in a microbiology class 2 biosafety cabinet, with a focused intensive rinse on the belly region. The sampling was performed by immediately and firmly swabbing a 3 cm^2^ linear belly area using a sterile flocked swab, which was then placed in 1 mL of liquid Amies storage media recipient (Copan ESwabs, Italy). No specimens were harmed during this work and all specimens were later released back into the aquaculture facility

From each sample, 100 µL of Amies media was placed on Tryptic Soy agar (TSA), Drigalski agar, Nutrient agar, and general use Chromatic Detection agar plates (MLT, Romania), and spread until dry using a sterile Drigalski spatula and a Petri dish turntable (Schuett-Biotec, Germany). The plates were incubated at 27° C and colonies were counted using a colony counter (Schuett-Biotec, Germany) after 24 and 48 hours of incubation, prior to strain isolations. With few exceptions, fish bacteria are inhibited by incubation temperatures above 30° C [19], therefore the 27° C incubation temperature was chosen as a realistic compromise for sturgeon surface microbiota growth, exceeding this threshold temperature enhances heat shock protein (hsp) expression in juvenile specimens, especially the protein folding and heat stress marker hsp70 [20].

### 2.3. Taxonomic identification

Final taxonomic identification was performed with a Matrix-Assisted Laser Desorption/Ionization Time-of-Flight (MALDI-TOF) Mass Spectrometer Biotyper (MALDI-TOF Microflex LT/SH, BrukerDaltonik, Germany). MSP 96 target polished steel plate (Bruker Daltonics, Germany) spots were loaded with isolated colonies, and α-Cyano-4-hydroxycinnamic acid (Bruker Daltonics, Germany) matrix was added to the stock solution (acetonitrile, trifluoracetic acid and distilled water). The SR Library Database was used for the identification (Database CD BT BTyp2.0-Sec-Library 1.0) with MBT Compass (v.4.1) software.

## 3. Results and Discussions

The bacterial load and diversity varied significantly among *A. gueldenstaedti* individuals, although they had belonged to the same brood.

During this study we isolated and identified 11 bacterial species, out of which 3 particularly adherent types, (Table 1) *Bacillus cereus, Plesiomonas shigelloides* and *Aeromonas* spp are notable. These are of high interest for public and veterinary health monitoring, and especially relevant for development of fish skin graft wound dressing technologies [21].

**Figure 1.**
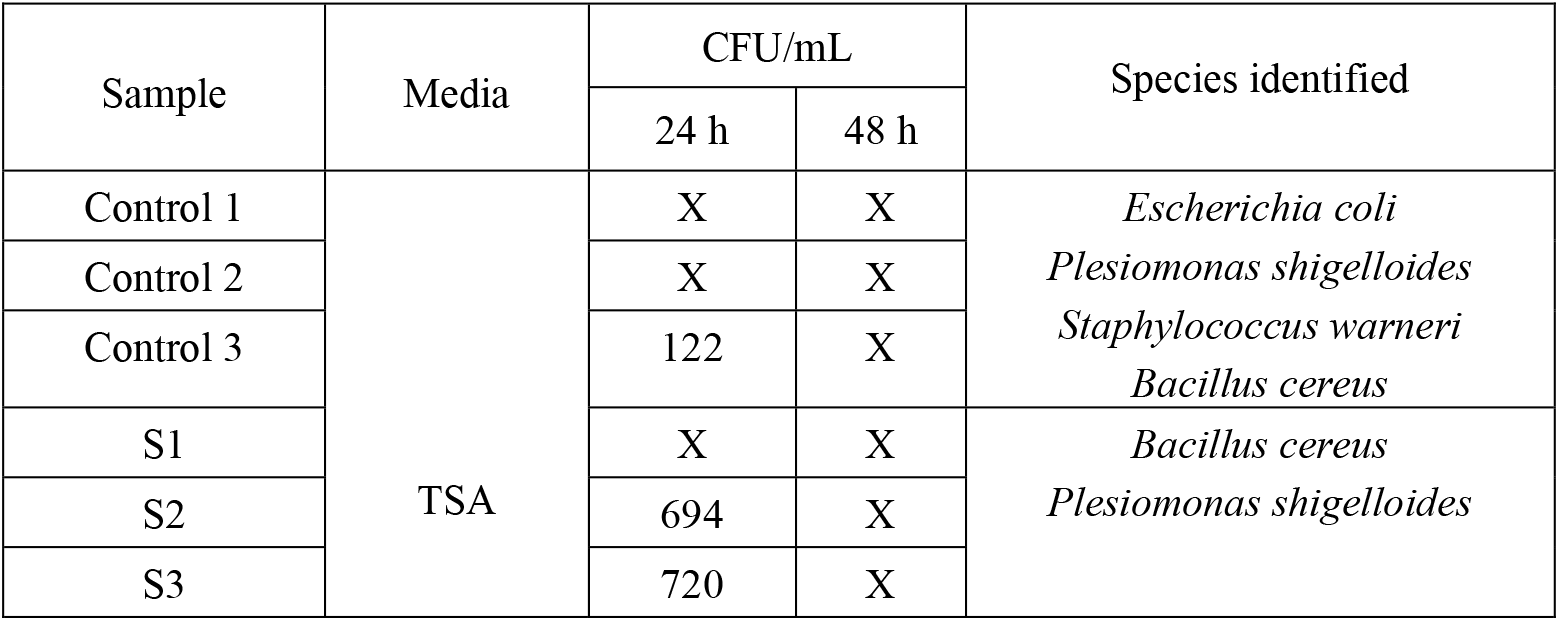
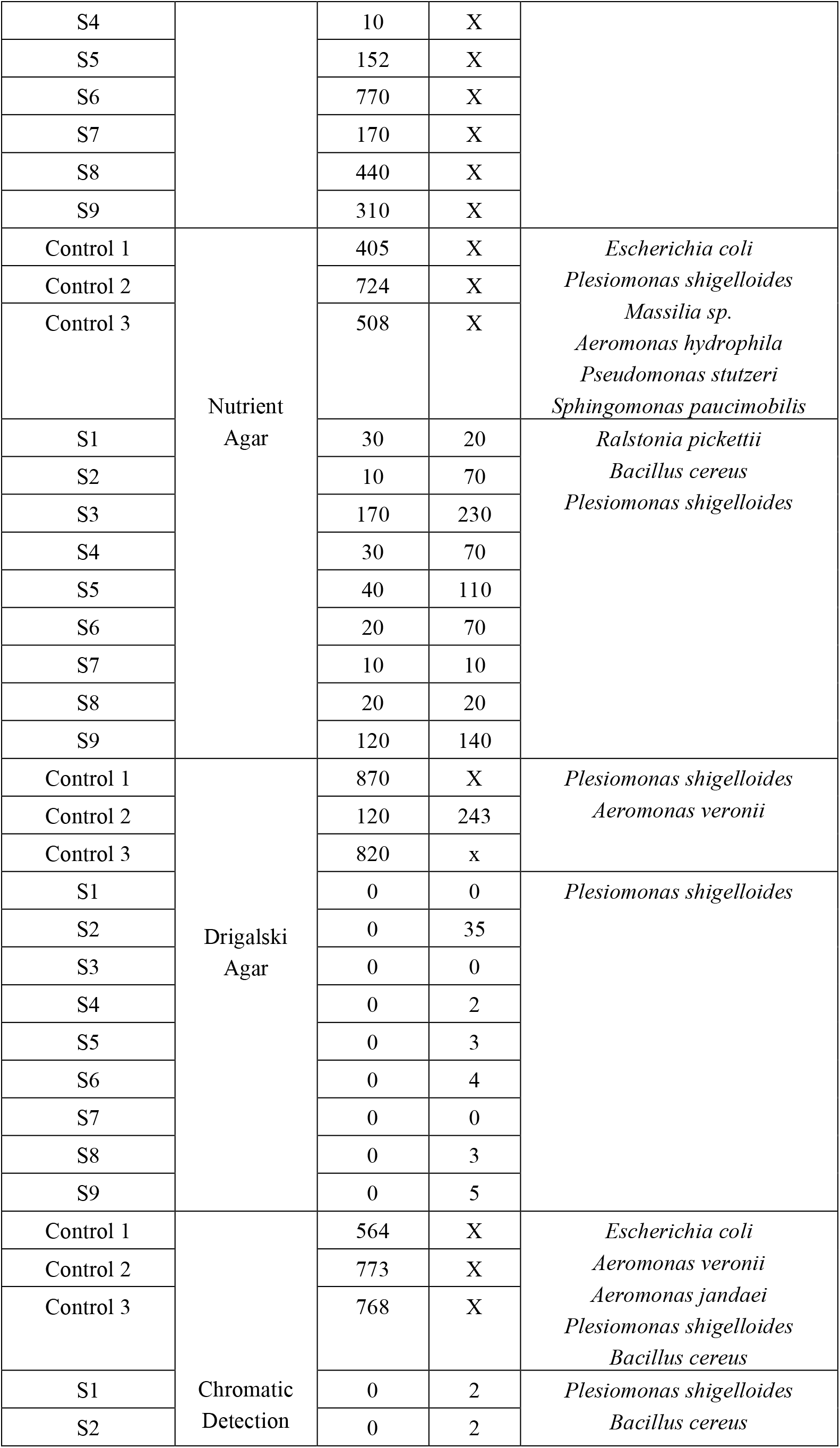
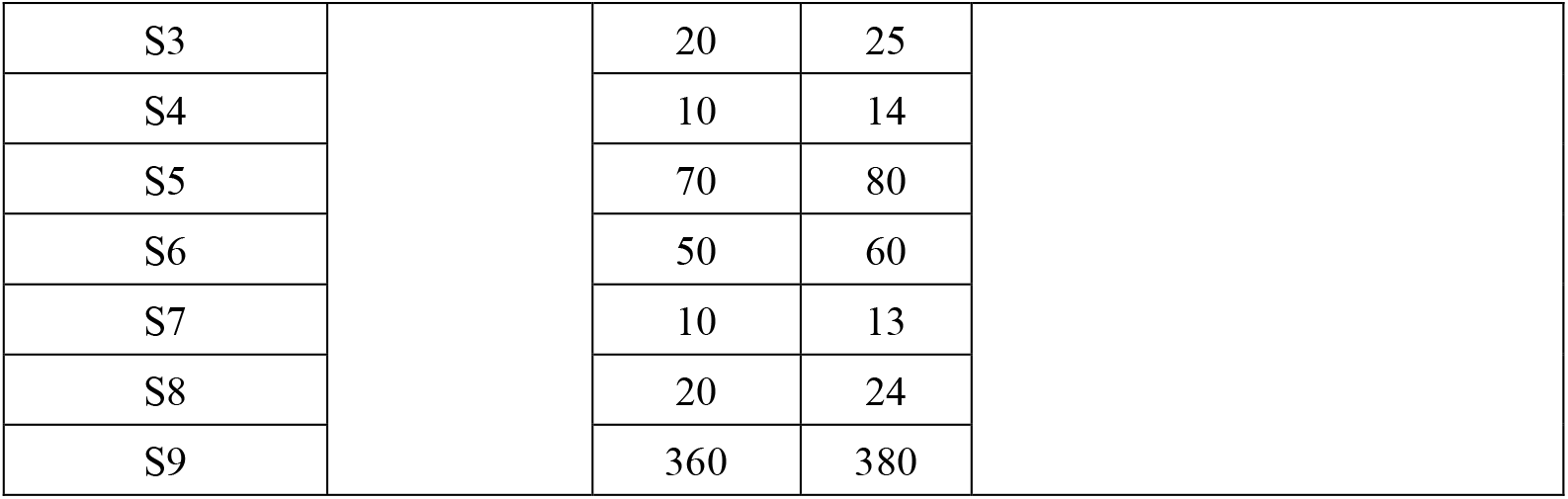
List of bacterial species identified from various culture media. X = Too Numerous To Count.

*Plesiomonas shigelloides* was the most prevalent species and grew on all media types that were used in this study, thus showing a great versatility and adaptability for which it’s already known [22]. This rod-shaped gram-negative pathogen discovered by Ferguson and Henderson in 1947 from clinical fecal samples [23] is native to aquatic organisms and has been isolated from fish, shellfish, crustaceans, marine mammals, amphibians, reptiles, and other aquatic lifeforms [24]. Although initially little attention was given to it as a potential pathogen, *P. shigelloides* now ranks with other established pathogens, like Aeromonas, as a potential cause of bacterial gastroenteritis [25–27]. However, unlike Aeromonas, how pH, temperature, turbidity, and conductivity affect *P. shigelloides* populations in freshwater ecosystems remain poorly understood [28]. The two taxa are known to be often found together, so much so that, traditionally, the genera Aeromonas and Plesiomonas have been taken together in the past [29, 30]. After three international workshops have described and accounted for many different aspects of these two genera they are now clearly distinguished as separate [29].

*Aeromonas* was the second most abundant genus, being represented by *A. hydrophila, A. veronii*, and *A. jandaei*, which all grew in the positive controlled groups, but none was found in rinsed groups, therefore indicating far less adhesivity than *P. shigelloides*. Bacterial adherence is considered a sign of higher pathogenicity [31–33], yet interestingly in fish, *P. shigelloides* infections are less virulent [34], and less frequent than those of Aeromonas sp. [22], although *P. shigelloides* was reported to adhere and enter eukaryotic intestinal host cells [35].

*Bacillus cereus*, another microorganism commonly associated with the aquatic environment [36, 37], was identified as being adherent on the tegument of the sturgeon specimens. There are reports of *Bacillus cereus* colonizing the intestinal tract of fishes [36, 38], and even being used as a biological agent against streptococcosis infections [38], however, to the best of our knowledge none have reported it as a skin pathogen in sturgeons. The gram positive rod-shaped bacteria is nevertheless a pathogen to humans, being transmitted via consumption of infected fish [39] and it’s also known for its versatility in terms of temperature, pH and is responsible for food-borne outbreaks [40-42].

*Pseudomonas stutzeri* and *Sphingomonas paucimobilis* were found growing only on the nutrient media and only on controls (probably non-adherent). The first is a denitrifying bacterium with ubiquitous environmental distribution, and has additionally been identified as an opportunistic human pathogen. The species has been suggested as an exemplary model organism for investigations into denitrification processes due to its unique metabolic capabilities. Moreover, numerous strains exhibit inherent natural transformation abilities and some possess the capacity for dinitrogen fixation [43]. The latter species, formerly known as *Pseudomonas paucimobilis* and CDC group IIk-1 [44], is widely distributed in water, including water sources in the hospital environment.

*Massilia aurea* was also identified as a possible non-adherent which grew only on nutrient media. The species is worth taking into account as it was first identified only almost two decades ago in drinking water in Spain [45]. There is little information regarding *Massilia aurea* to date, however Massilia strains are extensively distributed and perform diverse ecological roles. Initial investigations indicate that Massilia is the predominant species within constructed wetland ecosystems, yet the specifics of its species makeup and distribution in these wetlands remain uncertain [46]. Massilia spp. play a constructive role in environmental enhancement, for instance, *M. aromaticvorans* (ML15P13T) possesses the capacity to break down harmful gaseous organic air pollutants such as benzene, toluene, ethylbenzene, and xylene isomers [47].

The most particularly interesting finding in this study was the skin-adherent presence of *Ralstonia pickettii*.

*Ralstonia pickettii* is an emerging Gram-negative opportunistic pathogen [48]. Formerly called *Pseudomonas pickettii* [49] or *Burkholderia pickettii* [50], *R. pickettii* is a waterborne organism of low virulence and minor clinical importance [49] that, to the best of our knowledge, has not been reported in sturgeons. However, the species was identified in biofilm formation in plastic water piping where it resisted disinfection with numerous potent agents, showing a high resilience [51], and was also found in samples of wild and aquaculture fresh seafood [52]. The identification of this species is of particular importance as it can be considered an extremophile due to it’s high oligotrophic and other resistance abilities [53].

## 4. Conclusions

In this study surprisingly rarely studied bacterial species have been identified, and some found to be adherent on the sturgeon skin. In particular, *Ralstonia pickettii* is hereby reported for the first time. There are currently no known links between this species and wild or aquaculture sturgeons. The detected adherent and non-adherent species raise questions and concerns in terms of current sturgeon microbial communities and how climate changes will fluctuate such populations.

Future studies examining the possible presence of microbiota within the skin layers are required to support the present preliminary data. Further, bioprospecting microbiota of wild living sturgeons is recommended due to the current climate and ecological changes. These include the increasing levels of water contamination with older and emergent pollutants, all of which can induce not only immune system alterations of fishes, but also significant microbial community changes that can lead to detrimental consequences of unknown extent. Another important study should be on the development of the microbial adherents as the fish grow and develop. With clear differences in bacterial populations and diversity it might be important in fish health in aquaculture to understand which individuals might be hosts for pathogenic species, and their characteristics.

## Institutional Review Board Statement

Not applicable

## Data Availability Statement

Data available from the corresponding author upon reasonable request.

## Acknowledgments

N/A

## References

[1] D.C. Little, R. Newton, and M. Beveridge, “Aquaculture: a rapidly growing and significant source of sustainable food? Status, transitions and potential”. Proceedings of the Nutrition Society, 2016. 75(3): p. 274–286.

[2] M. Troell, et al., “Does aquaculture add resilience to the global food system?” Proceedings of the National Academy of Sciences, 2014. 111(37): p. 13257–13263.

[3] G. Roussow, “Some considerations concerning sturgeon spawning periodicity.” Journal of the Fisheries Board of Canada, 1957. 14(4): p. 553–572.

[4] H.J. Puglis, R.D. Calfee, and E.E. Little, “Behavioral effects of copper on larval white sturgeon”. Environmental Toxicology and Chemistry, 2019. 38(1): p. 132–144.

[5] B. Moshtaghi, et al., “Histopathological and bacterial study of Persian sturgeon fry, Acipenser persicus (Borodin, 1897) exposed to copper sulfate and potassium permanganate”. Journal of Parasitic Diseases, 2016. 40(3): p. 779–784.

[6] M. Santi, et al., “A survey of bacterial infections in sturgeon farming in Italy”. Journal of Applied Ichthyology, 2019. 35(1): p. 275–282.

[7] M.K. Dias, et al., “Lethal dose and clinical signs of Aeromonas hydrophila in Arapaima gigas (Arapaimidae), the giant fish from Amazon”. Veterinary microbiology, 2016. 188: p. 12–15.

[8] A. Choudhury, and T.A. Dick, “Parasites of lake sturgeon, Acipenser fulvescens (Chondrostei: Acipenseridae), from central Canada”. Journal of Fish Biology, 1993. 42(4): p. 571–584.

[9] M.R. Noei, “Parasitic worms of Acipenser stellatus, A. gueldenstaedtii, A. nudiventris and Huso huso (Chondrostei: Acipenseridae) from the southwest shores of the Caspian Sea”. 2011.

[10] O. Bauer, O. Pugachev, and V. Voronin, “Study of parasites and diseases of sturgeons in Russia: a review”. Journal of Applied Ichthyology, 2002. 18(4-6): p. 420–429.

[11] A. Leprévost, et al., “Vertebral Development and Ossification in the Siberian Sturgeon (Acipenser Baerii), with New Insights on Bone Histology and Ultrastructure of Vertebral Elements and Scutes”. The Anatomical Record, 2017. 300(3): p. 437–449.

[12] C. Höhne, et al., “The immune system of sturgeons and paddlefish (Acipenseriformes): a review with new data from a chromosome-scale sturgeon genome”. Reviews in Aquaculture, 2021. 13(3): p. 1709–1729.

[13] I. Salinas, “The Mucosal Immune System of Teleost Fish”. Biology, 2015. 4(3): p. 525–539.

[14] S. Yang, et al., “The Structure of the Skin, Types and Distribution of Mucous Cell of Yangtze Sturgeon (Acipenser dabryanus)”. International Journal of Morphology, 2019. 37: p. 541–547.

[15] G.F. Weisel, “The Integument and Caudal Filament of the Shovelnose Sturgeon, Scaphirhynchus platorynchus”. The American Midland Naturalist, 1978. 100(1): p. 179–189.

[16] V.S. Raj, et al., “Skin mucus of Cyprinus carpio inhibits cyprinid herpesvirus 3 binding to epidermal cells”. Vet Res, 2011. 42(1): p. 92.

[17] S. Dash, et al., “Epidermal mucus, a major determinant in fish health: a review”. Iran J Vet Res, 2018. 19(2): p. 72–81.

[18] B. Tørud, and T. Håstein, “Skin lesions in fish: causes and solutions”. Acta Veterinaria Scandinavica, 2008. 50(1): p. S7.

[19] C.E. Starliper, “General and specialized media routinely employed for primary isolation of bacterial pathogens of fishes”. J Wildl Dis, 2008. 44(1): p. 121–32.

[20] H. Chen, et al., “Effects of Temperature on the Growth Performance, Biochemical Indexes and Growth and Development-Related Genes Expression of Juvenile Hybrid Sturgeon (Acipenser baerii & Acipenser schrenckii)”. Water, 2022. 14(15): p. 2368.

[21] M. Ibrahim, et al., “Fish Skin Grafts Versus Alternative Wound Dressings in Wound Care: A Systematic Review of the Literature”. Cureus, 2023. 15(3): p. e36348.

[22] C.J. Grim, Chapter 14 - “Aeromonas and Plesiomonas, in Foodborne Infections and Intoxications (Fourth Edition)”, J.G. Morris and M.E. Potter, Editors. 2013, Academic Press: San Diego. p. 229–237.

[23] W. Ferguson, and N. Henderson, “Description of strain C27: a motile organism with the major antigen of Shigella sonnei phase I”. Journal of bacteriology, 1947. 54(2): p. 179–181.

[24] T.D. Jagger, “Plesiomonas shigelloides-a veterinary perspective”. Infect Dis Rev, 2000. 2: p. 199–210.

[25] S.D. Holmberg, and J. Farmer III, “Aeromonas hydrophila and Plesiomonas shigelloides as causes of intestinal infections”. Reviews of infectious diseases, 1984. 6(5): p. 633–639.

[26] N. Khardori, and V. Fainstein, “Aeromonas and Plesiomonas as etiological agents”. Annual Reviews in Microbiology, 1988. 42(1): p. 395–419.

[27] D.A. Sack, et al., “Epidemiology of Aeromonas and Plesiomonas diarrhoea”. Journal of Diarrhoeal Diseases Research, 1988: p. 107–112.

[28] J.M. Janda, S.L. Abbott, and C.J. McIver, “Plesiomonas shigelloides Revisited”. Clin Microbiol Rev, 2016. 29(2): p. 349–74.

[29] J.J. Farmer, M.J. Arduino, and F.W. Hickman-Brenner, “The Genera Aeromonas and Plesiomonas, in The Prokaryotes: A Handbook on the Biology of Bacteria Volume 6: Proteobacteria: Gamma Subclass”, M. Dworkin, et al., Editors. 2006, Springer New York: New York, NY. p. 564–596.

[30] A. von Graevenitz, and C. Bucher, “Evaluation of differential and selective media for isolation of Aeromonas and Plesiomonas spp. from human feces”. Journal of Clinical Microbiology, 1983. 17(1): p. 16–21.

[31] M.S. Donnenberg, and J.P. Nataro, “Methods for studying adhesion of diarrheagenic Escherichia coli”, in Methods in Enzymology. 1995, Academic Press. p. 324–336.

[32] J. Pizarro-Cerdá, and P. Cossart, “Bacterial Adhesion and Entry into Host Cells”. Cell, 2006. 124(4): p. 715–727.

[33] H.H. Niemann, W.-D. Schubert, and D.W. Heinz, “Adhesins and invasins of pathogenic bacteria: a structural view”. Microbes and Infection, 2004. 6(1): p. 101–112.

[34] L. Ashkenazi-Hoffnung, and S. Ashkenazi, “143 - Plesiomonas shigelloides”, in Principles and Practice of Pediatric Infectious Diseases (Sixth Edition), S.S. Long, Editor. 2023, Elsevier: Philadelphia. p. 850-851.e1.

[35] C. Theodoropoulos, et al., “Plesiomonas shigelloides enters polarized human intestinal Caco-2 cells in an in vitro model system”. Infect Immun, 2001. 69(4): p. 2260–9.

[36] X. Ke, et al., “A Bacillus cereus NY5 strain from tilapia intestine antagonizes pathogenic Streptococcus agalactiae growth and adhesion in vitro and in vivo”. Aquaculture, 2022. 561: p. 738729.

[37] G. Chandra, I. Bhattacharjee, and S. Chatterjee, “Bacillus cereus infection in stinging catfish, Heteropneustes fossilis (Siluriformes: Heteropneustidae) and their recovery by Argemone mexicana seed extract”. 2015.

[38] M. Wang, et al., “Effect of Bacillus cereus as a water or feed additive on the gut microbiota and immunological parameters of Nile tilapia”. Aquaculture Research, 2017. 48(6): p. 3163–3173.

[39] A. Doménech-Sánchez, et al., “Emetic disease caused by Bacillus cereus after consumption of tuna fish in a beach club”. Foodborne Pathog Dis, 2011. 8(7): p. 835–7.

[40] R. Labbé, and T. Rahmati, “Growth of Enterotoxigenic Bacillus cereus on Salmon (Oncorhynchus nerka)”. Journal of Food Protection, 2012. 75(6): p. 1153–1156.

[41] N. Jessberger, et al., “The Bacillus cereus Food Infection as Multifactorial Process”. Toxins, 2020. 12(11): p. 701.

[42] B. Glasset, et al., “Bacillus cereus-induced food-borne outbreaks in France, 2007 to 2014: epidemiology and genetic characterisation”. Eurosurveillance, 2016. 21(48): p. 30413.

[43] J. Lalucat, et al., “Biology of Pseudomonas stutzeri”. Microbiol Mol Biol Rev, 2006. 70(2): p. 510–47.

[44] J.P. Steinberg, and E.M. Burd, 238 - “Other Gram-Negative and Gram-Variable Bacilli”, in Mandell, Douglas, and Bennett’s Principles and Practice of Infectious Diseases (Eighth Edition), J.E. Bennett, R. Dolin, and M.J. Blaser, Editors. 2015, W.B. Saunders: Philadelphia. p. 2667-2683.e4.

[45] V. Gallego, et al., “Massilia aurea sp. nov., isolated from drinking water”. International Journal of Systematic and Evolutionary Microbiology, 2006. 56(10): p. 2449–2453.

[46] A. Xu, et al., “Dynamic distribution of Massilia spp. in sewage, substrate, plant rhizosphere/phyllosphere and air of constructed wetland ecosystem”. Frontiers in Microbiology, 2023. 14.

[47] J. Son, et al., “Massilia aromaticivorans sp. nov., a BTEX degrading bacterium isolated from Arctic soil”. Current Microbiology, 2021. 78: p. 2143–2150.

[48] M. Ryan, J. Pembroke, and C. Adley, “Ralstonia pickettii: a persistent gram-negative nosocomial infectious organism”. Journal of Hospital infection, 2006. 62(3): p. 278–284.

[49] D.A. Bruckner, and P. Colonna, “Nomenclature for aerobic and facultative bacteria”. Clinical infectious diseases, 1997: p. 1–10.

[50] E. Yabuuchi, et al., “Transfer of two Burkholderia and an Alcaligenes species to Ralstonia gen. nov.: proposal of Ralstonia pickettii (Ralston, Palleroni and Doudoroff 1973) comb. nov., Ralstonia solanacearum (Smith 1896) comb. nov. and Ralstonia eutropha (Davis 1969) comb. nov.”. Microbiology and immunology, 1995. 39(11): p. 897–904.

[51] R.L. Anderson, et al., “Effect of disinfectants on pseudomonads colonized on the interior surface of PVC pipes”. American Journal of Public Health, 1990. 80(1): p. 17–21.

[52] M. Boulares, et al., “Characterisation and identification of spoilage psychotrophic Gram-negative bacteria originating from Tunisian fresh fish”. Annals of Microbiology, 2013. 63(2): p. 733–744.

[53] C. Adley, M. Ryan, J. Pembroke, F.M. Saieb, “Ralstonia pickettii: biofilm formation in high-purity water”. Biofilms: persistence and ubiquity. Biofilm Club, Powys, 2005, pp 261–271

